# Current achievements and future developments of a novel AI based visual monitoring of beehives in ecotoxicology and for the monitoring of landscape structures

**DOI:** 10.1101/2020.02.04.933580

**Authors:** Frederic Tausch, Katharina Schmidt, Matthias Diehl

## Abstract

Honey bees are valuable bio-indicators. As such, they hold a vast potential to help shed light on the extent and interdependencies of factors influencing the decline in the number of insects. However, to date this potential has not yet been fully leveraged, as the production of reliable data requires large-scale study designs, which are very labour intensive and therefore costly.

A novel Artificial Intelligence (AI) based visual monitoring system could enable the partial automatization of data collection on activity, forager loss and impairment of the central nervous system. The possibility to extract features from image data could prospectively also allow an assessment of pollen intake and a differentiation of dead bees, drones and worker bees as well as other insects such as wasps or hornets.

The technology was validated in different studies with regards to its scalability and its ability to extract motion and feature related information.

The prospective possibilities were analyzed regarding their potential to enable advances both within ecotoxicological research and the monitoring of pollinator habitats.

## Abstract in German

Honigbienen sind nützliche Bio-Indikatoren. Sie besitzen ein großes Potenzial, Licht in das Ausmaß und die Wechselwirkungen der Faktoren zu bringen, die den Rückgang der Insektenzahl beeinflussen. Dieses Potenzial wurde bisher noch nicht vollständig ausgeschöpft, da die Produktion zuverlässiger Daten aufwendige und arbeitsintensive Studiendesigns erfordert, die mit hohen Kosten verbunden sind.

Ein neuartiges, auf künstlicher Intelligenz (KI) basierendes visuelles Monitoringsystem könnte eine teilweise Automatisierung der Datenerfassung über Aktivität, Sammelbienenverlust und Beeinträchtigungen des zentralen Nervensystems ermöglichen. Die Möglichkeit, Merkmale aus den Bilddaten zu extrahieren, könnte darüber hinaus zukünftig auch eine Bewertung des Polleneintrags und eine Unterscheidung von toten Bienen, Drohnen und Arbeitsbienen sowie anderen Insekten wie Wespen oder Hornissen ermöglichen. Die Technologie wurde in verschiedenen Studien hinsichtlich ihrer Skalierbarkeit und ihrer Fähigkeit, bewegungs- und merkmalsbezogene Informationen zu extrahieren, validiert.

Die Möglichkeiten wurden hinsichtlich ihres Potenzials analysiert, Fortschritte sowohl in der ökotoxikologischen Forschung als auch im Monitoring von Bestäuber-Lebensräumen zu ermöglichen.

## Introduction

Honey bee colonies can act as detectors for harmful substances by either signaling the existence of toxic molecules through high mortality rates or by accumulating residues for not acutely lethal substances e.g., of heavy metals, fungicides and herbicides in pollen, nectar or larvae (Celli, 1983; Porrini et al., 2002). They were first used as bio-indicators to monitor environmental quality in 1935 (Crane, 1984). The detection of pesticide use is one of the research fields in which bee monitoring has since been applied (Atkins et al., 1981; Celli, 1983; Mayer and Lunden, 1986; Mayer et al., 1987; Celli et al., 1988; Celli and Porrini, 1991; Celli et al., 1991; Porrini et al., 1996). With about a quarter of its inhabitants being active foragers, the condition of a colony mirrors the state of its habitat. Among the requisites which make the colonies especially suitable environmental indicators are that they can be easily held by beekeepers, that their foragers cover large areas and that they collect samples like pollen or nectar out of self-interest. (Celli and Maccagnani, 2003).

Honey bee colony development depends on many factors such as but not limited to queen age, nutrition, colony strength, pathogens and parasites as well as regional particularities. Therefore, large sample sizes are necessary in order to generate objective insights into causal relationships of hazards towards honey bees. Within the German bee project, which was aimed at understanding the causes of honey bee colony collapses, more than 1.200 bee hives in 125 places across the country were monitored between 2004 and 2009. The study brought many correlations to light but also left some questions open. The authors presume that a study design suitable to record sublethal or chronic effects might reveal a negative effect of pesticides regarding colony collapse which they were not able to detect. (Genersch et al., 2010).

Because large scale studies using bees as bioindicators are very time and labour intensive, their number is still quite small. In 1978, Giordani et al. were able to demonstrate the highly toxic effect of the chlorinated hydrocarbon insecticide Endosulfan. Yet, many years and several studies were needed to provide enough evidence to change a limitation of the use of the substance. Later, in a large-scale monitoring project in northern Italy, bee mortality was recorded for several hundred hives under both high and low chemical pressure from farming. By analyzing dead bees from hives with especially high numbers of casualties it was possible to identify the molecules responsible for 76 % of registered mass-deaths. Nevertheless, one shortcoming of the design the authors mentioned was that the number of dead bees collected was only a conservative estimate as the losses caused by lethal doses in the field could not be recorded. (Celli and Maccagnani, 2003).

These works demonstrate the potential of bee monitoring in various fields from pesticide regulation to general advances in research regarding bee health. However, they are pioneer projects and not representative for the way research is typically conducted. To date, factor analysis and preventive activities are built mostly on snapshot data from a small number of hives which can be collected more economically. The use of technology could help decrease the labour intensity and therefore the cost of such projects. A few systems, based on different technologies has recently been developed, yet still exhibit shortcomings.

There are counting systems, which try to quantify the incoming and outgoing bees at the entrance, e.g. BeeCheck with capacitive detection (Gombert et al., 2019). Due to their design, the counting systems only record the pollinators within a short distance. The information content of their sensory raw data is considerably reduced for evaluation with an imaging method. In complex situations, such as bees running on top of each other or group formation they are therefore prone to measurement inaccuracies and would consequently not be suitable for a robust assessment of mortality. With a visual system it is possible to follow each individual animal over a sequence of images. First scientific works could already present prototype systems, which used a camera system at the hive entrance to determine the parasite infestation (Schurischuster et al., 2018).

From 2017 to 2020, the EU-funded project IoBee aims to identify global changes in bee populations through the networking of data from bee colonies. The data collection within the project is supported by technical sensor systems. However, these partly specialized, partly integrated sensor systems detect only temperature, humidity, sound and weight to determine the health of individual hives. They are not designed to collect information about activity, forager loss or foraging intensity.

AI has a high potential to contribute to data in areas, which can only be collected from images (Bozek et al. 2018). Because Neural Networks, like all software are scalable and once trained able to extract results very precisely, their use could vastly improve the quantity and quality of information which can be gathered from the monitoring of bee hives.

## Materials and Methods

An AI based visual monitoring technology for bee hives has been developed (figure 1), in order to create a cost-efficient way to autonomously collect data on a larger scale with high quality. It combines hardware and software components. A camera device is attached to the hive entrance to record all bees entering and leaving. As there are usually no energy and network connections available at bee hives during field studies, the device was equipped with a solar panel. A UMTS-connection makes sure the software on the device can be accessed and updated remotely. Different software modules subsequently analyze the image data both on-device and with the help of cloud computing resources. Deep learning algorithms are used which have been trained through exposure to selected exemplary data sets. These methods of AI enable the collection of objective data and still allow a verification of the results at a later time, because image segments with individual bees and video sections are simultaneously and sequentially archived in a database. The technology can be applied to generate both motion and feature related insights.

**Fig. 1.**
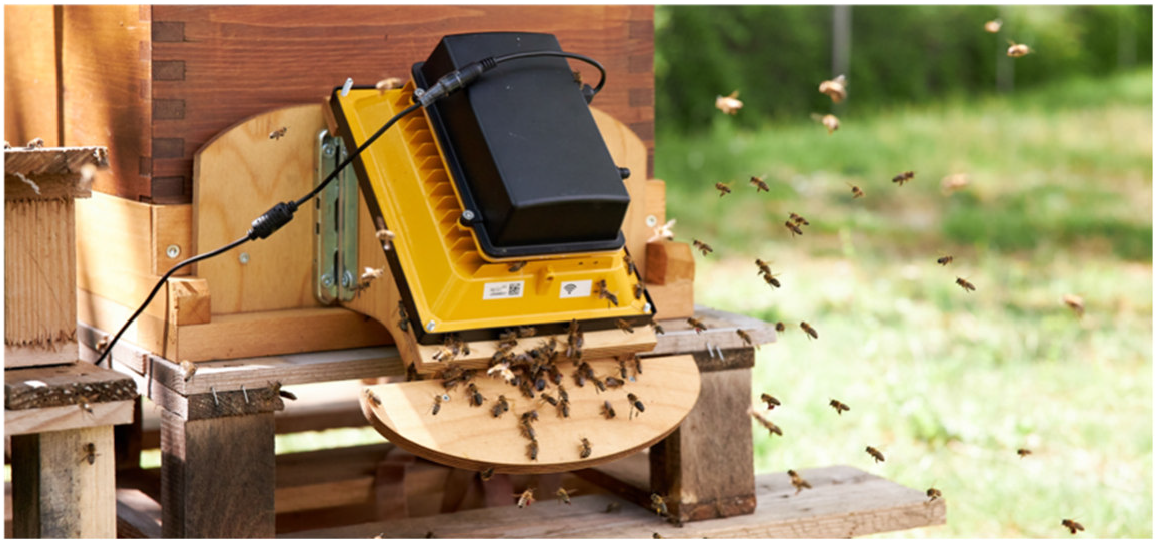
Visual monitoring device in front of the hive entrence Visuelles Monitoring-System vor dem Eingang des Bienenstocks

### Motion-related analysis

The ingress and egress of bees is captured on camera at the hive entrance. Neural Networks detect each bee and track its movement while it passes through the camera’s field of view, which measures about 145 x 108 mm^2^ (figure 2, A). Subsequently, different aspects can be analyzed:

- Level of activity derived from number of incomings and outgoings
- Loss as the proportion of outgoings which do not return
- Motion patterns of individual bees within the cameras window frame

**Fig. 2.**
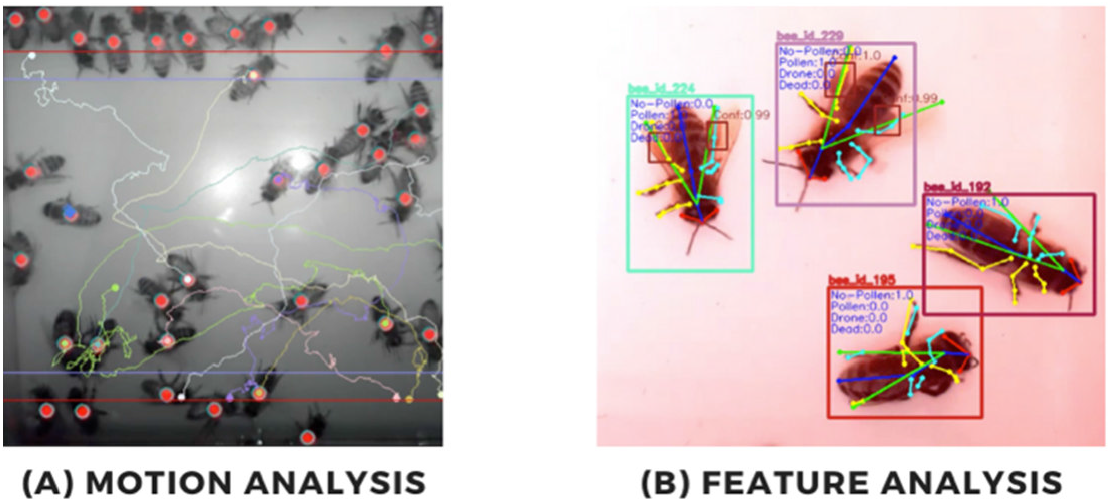
Visualiserung der Bewegungsanalyse (A) und Visualisierung der Featureanalyse (B) Visualisation of the motion Analysis (A) and visualization of the feature analysis (B)

### Feature-related analysis

Cropped single-entity images of bees entering and leaving their hive are uploaded into a cloud (figure 2, B). The collected data is processed by a multi modal neuronal network to perform the the following tasks:

- Recognition of pollen on bee’s legs
- Detection of whether a bee is dead or alive
- Differentiation of drones and workers
- Differentiation of other genera like wasps or hornets

In 2019, the monitoring system was tested in two test settings to validate different aspects of both the hardware and software components within the technology.

The practical scalability of the technology was tested as a precondition to apply it under realistic circumstances. The camera devices were attached to a total of 33 colonies in 14 locations in and around the city of Karlsruhe, at hives of local bee keepers, public institutions or companies.

In a different setting, bee activity was measured to detect changes in flight activity following neonicotinoid exposure. Within an Oomen feeding study in an agricultural area, the applicability of using the visual bee’s activity in pollinator risk assessment was evaluated. The study assessed the impact of the feeding with a neonicotinoid on daily activity and colony development. Eight hives were monitored, of which four were fed with 500 g of a sugar solution, including a concentration of 200 μg imidacloprid/kg of sugar solution, for ten consecutive days. The control group was fed the same amount of sugar solution during that exposure period (Gonsior et al., 2019).

In a separate proof-of-concept study, the possibilities of machine learning algorithms were explored to perform localization, classification and pose estimation tasks on video recordings (Marstaller et. al., 2019).

## Results

The results of the 2019 studies show first positive achievements for future application in the fields of ecotoxicology and for the monitoring of landscapes.

For both usecases, it is essential to build a scalable and failsafe system. Special methods for failure detection were developed, to ensure the uptime of the camera devices. Cloud monitoring alerts were set up for notification in case of failures to reduce downtime. Adding these mechanisms on different system levels made the devices independent and self-sufficient. This will make it possible in the future to continuously monitor large areas and remote locations.

Regarding the results of the Oomen study, figure 3 shows the change in activity per hive during the exposure period. The negative values represent bees leaving their hive while positive values represent bees returning. Values are plotted as the sum of bees per hour. During the exposure period, from the third day of exposure onwards the hives fed with the neonicotinoid displayed a significantly decreased level of activity. It has thus been demonstrated, the technology used can collect relevant parameters for ecotoxicological reseasrch, which could not be assessed before.

**Fig. 3.**
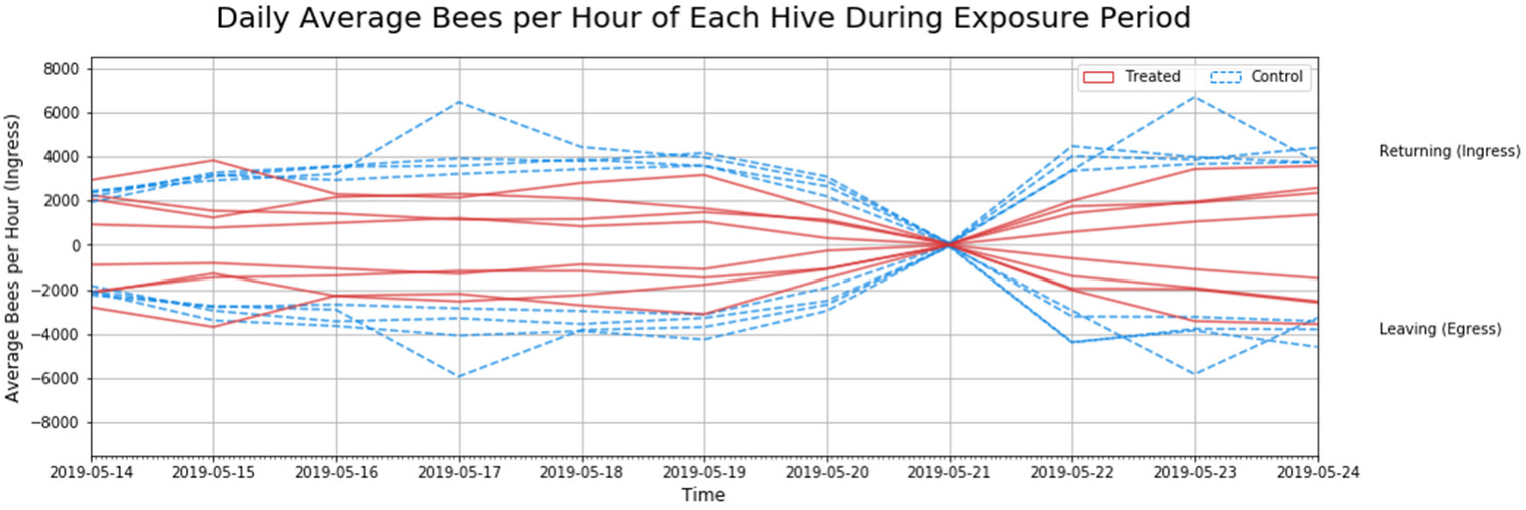
Aktivität am Bienenstockeingang nach dem Kontakt mit Imidacloprid (Gonsior et al., 2019) Activity at the hive entrance following contact with imidacloprid (Gonsior et al., 2019)

The detection of whether individual animals were drones, worker bees or different genuses like wasps and whether they were dead or alive could be achieved with the use of a cloud based multi network. It was also possible to detect whether bees carried pollen or not with a certain probability. Additionally, information on the bee’s pose could be recognized, including detailed information on the movement. This pilot work suggests that generating further insights beyond the activity of bees is feasible, which amplifies the advantages of using bees as bio-indicators.

## Conclusion

In first tests, AI based visual monitoring of bee hives has shown great potential to reliably capture and analyze honey bee’s motion and features. With the help of Neural Networks honey bees can be used as bio-indicators in new ways. Information about environmental factors can be collected, which has not yet been accessible.

A test-field with a networked system of prototypes for real-time analysis is in place in Karlsruhe for further testing. Algorithms are currently developed to be deployed in 2020 using this infrastructure. Data on activity, forager loss and food availability will be systematically collected. For this purpose, beehives will be monitored in rural and urban landscapes all over Baden-Württemberg. The aim of the project is to identify local and seasonal problems and evaluate landscape suitability as a habitat for insects. As the camera devices in the field can be updated remotely, software improvements can be distributed whenever algorithms are improved. The accuracy of the algorithms depends on the training data within the database. Therefore, it can be improved either by providing additional data sets or by improving the existing input data through data annotation and quality assurance by experts.

It has been proven feasible to set up a connected network of bee hives to remotely monitor hive activity. Tests of the scalability of the technology give reason to believe that it can be used to constantly and simultaneously monitor numerous hives. By allowing the monitoring of large numbers of colonies with minimal human labour, AI based visual monitoring of bee hives could therefore become a valuable tool in ecotoxicological field studies. It could in the future enable significantly larger study designs and therefore facilitate more reliable results. Once this it possible, the technology could be used to classify the magnitude of the detrimental effects of plant protection products as well as other environmental hazards on colonies.

In a collaborative study with Eurofins Agroscience Services Ecotox it has already been possible to quantitatively track bee activities. Improvements are currently in progress to achieve a level of accuracy at which an accurate determination of the precise loss of foragers can be derived from the quantitative ingress and egress activity. It could thus become possible for the first time to generate precise data of the loss of bees which died outside their hive because of lethal environmental effects. The data stream would be constant and could be collected without human action or assessment bias.

In addition to information about the level of activity and the precise loss of foragers, the motion profiles of each bee could be analyzed as an indicator of sub-lethal or chronic effects. Changes in motion patterns could indicate an impairment of the central nervous system or health related issues of colonies such as the deformed wing virus.

Furthermore, a quantitative assessment of pollen intake could be used to assess the extent of foraging activity. Low pollen intake at a number of neighbouring hives might indicate a temporal shortage in food supply within the respective region. This information could be utilized to implement targeted measures like the cultivation of plants which bloom during that period. As a result, the food situation would be improved not only for the honey bees but also for other pollinators. In this case the honey bee colonies could serve as a bio-indicator to identify environments in which there would not be sufficient food available to feed pollinators all year long. If merged with further information, such data could also contribute to ecological impact statements.

